# MGDrivE: A modular simulation framework for the spread of gene drives through spatially-explicit mosquito populations

**DOI:** 10.1101/350488

**Authors:** Héctor M. Sánchez C., Sean L. Wu, Jared B. Bennett, John M. Marshall

**Affiliations:** Division of Epidemiology and Biostatistics, School of Public Health, University of California, Berkeley, California, United States; Biophysics Graduate Group, Division of Biological Sciences, College of Letters and Science, University of California, Berkeley, California, United States; Innovative Genomics Institute, Berkeley, California, United States

## Abstract

Malaria, dengue, Zika, and other mosquito-borne diseases continue to pose a major global health burden through much of the world, despite the widespread distribution of insecticide-based tools and antimalarial drugs. The advent of CRISPR/Cas9-based gene editing and its demonstrated ability to streamline the development of gene drive systems has reignited interest in the application of this technology to the control of mosquitoes and the diseases they transmit. The versatility of this technology has also enabled a wide range of gene drive architectures to be realized, creating a need for their population-level and spatial dynamics to be explored. To this end, we present MGDrivE (Mosquito Gene Drive Explorer): a simulation framework designed to investigate the population dynamics of a variety of gene drive architectures and their spread through spatially-explicit mosquito populations. A key strength of the MGDrivE framework is its modularity: a) a genetic inheritance module accommodates the dynamics of gene drive systems displaying user-defined inheritance patterns, b) a population dynamic module accommodates the life history of a variety of mosquito disease vectors and insect agricultural pest species, and c) a landscape module accommodates the distribution of insect metapopulations connected by migration in space. Example MGDrivE simulations are presented to demonstrate the application of the framework to CRISPR/Cas9-based homing gene drive for: a) driving a disease-refractory gene into a population (i.e. population replacement), and b) disrupting a gene required for female fertility (i.e. population suppression), incorporating homing-resistant alleles in both cases. We compare MGDrivE with other genetic simulation packages, and conclude with a discussion of future directions in gene drive modeling.

## Introduction

The advent of CRISPR/Cas9-based gene editing technology and its application to the engineering of gene drive systems has led to renewed excitement in the use of genetics-based strategies to control mosquito vectors of human diseases and insect agricultural pests [1-3]. Applications to control mosquito-borne diseases have gained the most attention due to the major global health burden they pose through much of the world and the difficulty of controlling them using currently-available tools. For malaria, recent declines in transmission have been seen following the wide-scale distribution of bed nets and antimalarial drugs [4]; however, model-based projections suggest that additional tools will be required to eliminate the disease from highly-endemic areas [5]. For dengue, there is also a need for novel vector control strategies, as the disease is rising in global prevalence and there is currently no cure or vaccine available that is effective against all four serotypes [6]. The recent demonstration of a CRISPR-based gene drive system in Drosophila [7], followed months later by a Zika outbreak in Brazil [8], has prompted development of gene drive technology for *Aedes aegypti*, the primary mosquito vector of Zika, dengue, and Chikungunya, as well as broad development targeting other mosquito species, such as the Anophelines which transmit malaria.

The ease of gene editing afforded by the discovery of CRISPR has also led to significant versatility in terms of the gene drive systems that are now realizable [3, 9]. Prior to the advent of CRISPR, homing endonuclease genes (HEGs) were envisioned to cleave a specific target site lacking the HEG and to be copied to this site by serving as a template for homology-directed repair (HDR), effectively converting a heterozygote into a homozygote and biasing inheritance in favor of the HEG [10]. These dynamics have been demonstrated for a HEG targeting a synthetic target site in the main African malaria vector, *Anopheles gambiae* [11], and steps have also been taken towards engineering an alternative approach in which the HEG is located on the Y chromosome and cleaves the X chromosome in multiple locations, biasing inheritance in its favor as it induces an increasingly male sex bias in the population [12]. A vast range of additional approaches for biasing inheritance are now being proposed, including several threshold-dependent systems that may permit confineable and reversible releases [13-15], and remediation systems that could be used to remove effector genes and possibly entire drive systems from the environment in the event of unwanted consequences [16]. For instance, an ERACR system (Element for the Reversal of the Autocatalytic Chain Reaction) has been proposed that consists of a homing system with a target site corresponding to the original drive system, essentially removing the original drive as it homes into it, and utilizing the Cas9 of the first drive thus also removing this through the homing process [17, 18].

Understanding how these systems are expected to behave in real ecosystems requires a flexible modeling framework that can accommodate a range of inheritance patterns, specific details of the species into which the constructs are to be introduced, and details of the landscape through which spatial spread would occur. To this end, we present MGDrivE (Mosquito Gene Drive Explorer): a flexible simulation framework designed to investigate the population dynamics of a variety of gene drive systems and their spread through spatially-explicit populations of mosquito species and other insect species. A key strength of the MGDrivE framework is its modularity. A genetic inheritance module allows the inheritance dynamics of a wide variety of drive systems to be accommodated. An independent population dynamic module allows the life history of a variety of mosquito disease vectors and insect agricultural pests to be accommodated. Thirdly, a landscape module accommodates the distribution of insect metapopulations in space, with movement through the resulting network determined by dispersal kernels. The model can be run in either a deterministic or stochastic form, allowing the chance events that occur at low population or genotype frequencies to be simulated.

What separates MGDrivE from other gene drive modeling frameworks is its ability to simulate a wide array of user-specified inheritance-modifying systems at the population level within a single, computationally efficient framework that also incorporates mosquito life history and landscape ecology. Other frameworks exist that have been designed for more general purposes and applied to specific questions related to gene drive (Table 1) – for instance, Eckhoff *et al.* [19] used the EMOD malaria model to simulate the spread of homing-based gene drive systems through spatial populations of *An. gambiae*. EMOD is open source and a powerful modeling framework; but significant effort is required from users to redefine genetic control strategies, mosquito life history parameters and landscape details. Magori *et al.* [20] created Skeeter Buster by extending the CIMSiM (container-inhabiting mosquitoes simulation model) model [21] to incorporate genetic inheritance and spatial structure. The Skeeter Buster framework captures the most pertinent mosquito ecology considerations, but is not open source and can only simulate a handful of genetic control strategies [22]. The SLiM genetic simulation framework [23] is capable of modeling the spread of a large variety of user-defined gene drive systems through metapopulations; however, it is not currently capable of accommodating life history ecology and overlapping generations.

**Table 1.** Comparison of spatially-explicit gene drive models.

|  | Inheritance Patterns | Life History Ecology | Spatial and landscape details | Software |
| --- | --- | --- | --- | --- |
| MGDrive | Very flexible, can be user-specified | Egg-larva-pupa-adult, density-dependence at larval stage, not responsive to environmental variables at present | Metapopulations distributed in space, connected by migration | R package, open source |
| EMOD [19] | Homing-based gene drive, could be extended with effort | Egg-larva-pupa-adult, density-dependence at larval stage, responsive to environmental variables | Populations arranged on a grid, each representing 1 km<sup>2</sup>, connected by migration | Java Script Open Notation (JSON) feeds into executable file, open source |
| Skeeter Buster [22] | Homing-based gene drive, release of insects carrying a conditional lethal, etc., cannot be user-specified | Egg-larva-pupa-adult, density-dependence at larval stage, responsive to environmental variables | Households and containers modeled explicitly, connected by migration | Executable file, not open source |
| SLiM [23] | Very flexible, can be user-specified | Discrete generations, no life history at present | Can model either connected metapopulations or cells on a grid | Scripting environment with graphical user interface, open source |

In this paper, we describe the key components of the MGDrivE framework – namely, the genetic inheritance, mosquito life history and landscape/metapopulation modules. We then provide a demonstration of the application of the framework to CRISPR-based homing gene drive systems for: a) driving a disease-refractory gene into a population (i.e. population replacement), and b) disrupting a gene required for female fertility (i.e. population suppression), incorporating homing-resistant alleles. We conclude with a discussion of future applications of genetic simulation packages in the field of gene drive modeling.

## Design and Implementation

The MGDrivE framework is a genetic and spatial extension of the lumped age-class model of mosquito ecology [24] modified and applied by Deredec *et al.* [25] to the spread of homing gene drive systems, and by Marshall *et al.* [26] to population-suppressing homing systems in the presence of resistant alleles. The framework incorporates the egg, larval, pupal and adult life stages, with egg genotypes determined by maternal and paternal genotypes and the allelic inheritance pattern. In MGDrivE, by treating the lumped age-class model equations in a variable-dimension tensor algebraic form, the population dynamic equations can be left unchanged while modifying the dimensionality of the tensor describing inheritance patterns, as required by the number of genotypes associated with the drive system. Spatial dynamics are then accommodated through a metapopulation structure in which lumped age-class models run in parallel and migrants are exchanged between metapopulations at defined rates. These operations are accommodated by the tensor modeling framework, and full details of this framework are provided in the S1 File.

The core simulation framework is being developed in R (https://www.r-project.org/) with certain routines in Rcpp for computational speed. By combining the tensor modeling framework with object-oriented programming, the genetic, life history and spatial components of the model are able to be separated into “modules” to facilitate ease of modification. Within this architecture, each module may be conveniently altered independently of the others. For instance: a) a range of gene drive systems may be explored for a given mosquito species in a given landscape, b) one species may be substituted for another, provided its sequence of life history events is comparable, and c) gene drive spread may be modeled through a range of landscapes, while leaving the rest of the model untouched. We now describe the three distinct modules of the MGDrivE framework – inheritance, life history and spatial structure – in more detail.

## Modules

### 1. Genetic Inheritance

The fundamental module for modeling gene drive dynamics is that describing genetic inheritance. In MGDrivE, this is embodied by a three-dimensional tensor referred to as an “inheritance cube” (Figure 1). Each gene drive system has a unique R file containing the three-dimensional inheritance cube. The first and second dimensions of the inheritance cube refer to the maternal and paternal genotypes, respectively, and the third dimension refers to the offspring genotype. The cube entries for each combination of parental genotypes represent the proportion of offspring that are expected to have each genotype, and should sum to one, as fitness and viability are accommodated separately.

**Fig 1.**
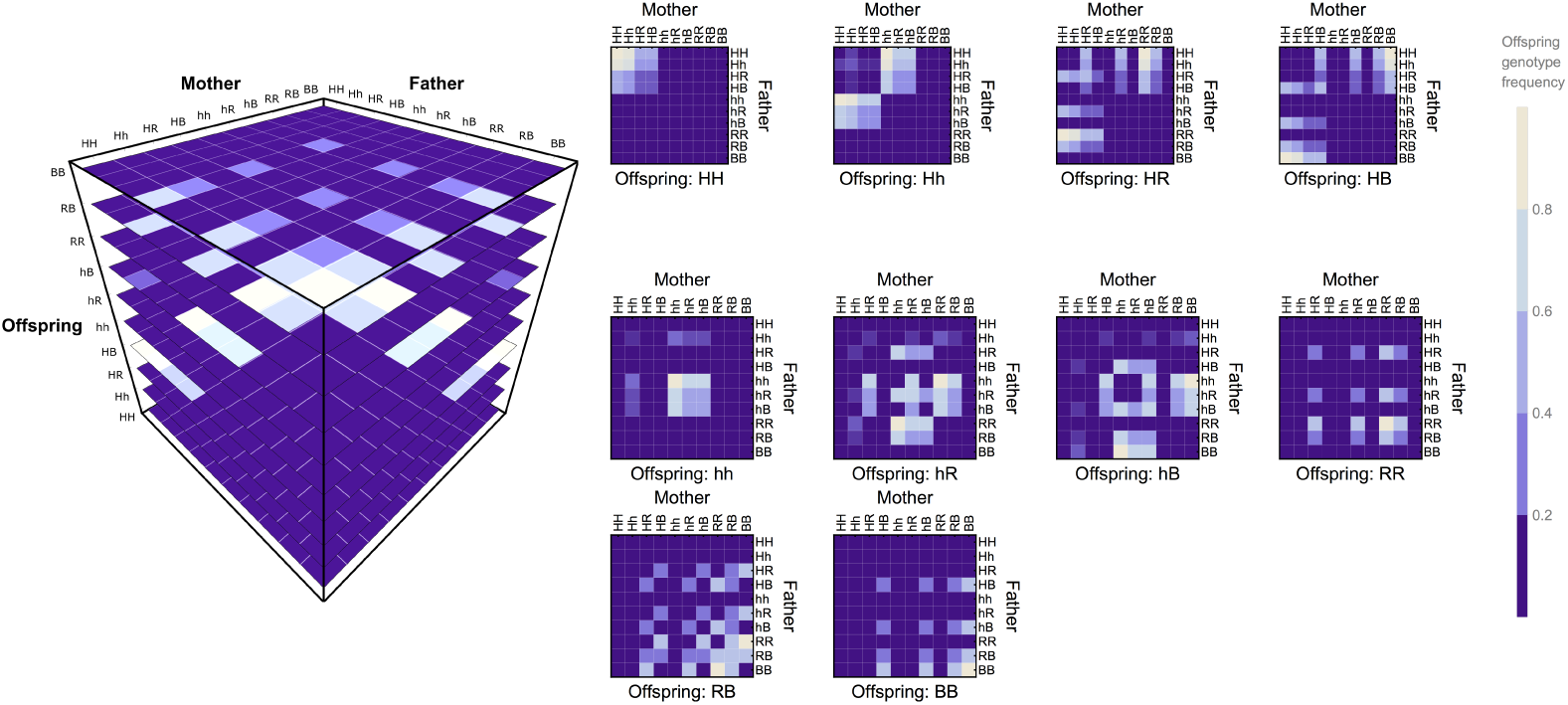
Inheritance module. Genetic inheritance is embodied by a three-dimensional tensor referred to as an inheritance cube (left), here depicted for a CRISPR/Cas9-based homing construct. Maternal and paternal genotypes are depicted on the x and y-axes and offspring genotypes on the z-axis, with slices of the cube pertaining to each offspring genotype shown to the right. The inheritance pattern shown deviates from standard Mendelian inheritance such that, in the germline of Hh parents, the majority of wild-type (h) alleles are converted into homing (H) alleles, while a small proportion are converted into in-frame resistant (R) and out-of-frame resistant alleles (B). For the example pictured, the frequency of accurate homing given cleavage in Hh heterozygotes is 98%, with the remaining 2% of wild-type alleles being converted to either in-frame (1%), or out-of-frame (1%) resistant alleles. Offspring genotype frequencies for each parental cross are depicted according to the shading scale (right).

The R function that builds the inheritance cube may receive a number of user-defined input parameters. For a homing-based drive system, for instance, the list of input parameters should include the homing efficiency, the rate of in-frame resistant allele generation and the rate of out-of-frame or otherwise costly resistant allele generation [26-28]. In-frame resistant alleles are those for which the coding frame of the target site is not altered, leading to minimal fitness effects, while out-of-frame resistant alleles disrupt the coding frame and hence function of the target site, leading to significant fitness effects. These parameter values should be used to populate the entries of the inheritance cube. Input parameters also include those associated with organisms having each genotype – for instance, genotype-specific: a) fertility rates, b) male mating fitness, c) sex bias at emergence, d) adult survival rates, and e) male and female pupatory success. These parameters feed into the mosquito life history module, that will be described next, and modify the tensor equations in that module in order to produce the desired biological effect. Finally, a “viability mask” is applied to the offspring genotypes to remove unviable genotypes from the population.

At the time of publication, the MGDrivE package includes inheritance cubes for: a) standard Mendelian inheritance, b) homing-based drive intended for population replacement or suppression [26, 27, 29, 30], c) *Medea* (a maternal toxin linked to a zygotic antidote) [31], d) other toxin-antidote-based underdominant systems such as UD^MEL^ [13, 15, 32], e) reciprocal chromosomal translocations [14, 33], f) *Wolbachia* [34], and g) the RIDL system [35] (release of insects carrying a dominant lethal gene). Details of each of these systems are provided in the user documentaion at https://marshalllab.github.io/MGDrivE/docs/reference/.

### 2. Mosquito Life History

The mosquito life history module follows from the lumped age-class model of Hancock and Godfray [24] adapted by Deredec *et al.* [25]. In this model (depicted in Figure 2), the insect life cycle is divided into four stages – egg (E), larva (L), pupa (P) and adult (M for male and F for female). In MGDrivE, each life stage is associated with a genotype. Adult females mate once and produce batches of eggs from the sperm of the same male, so they obtain a composite genotype upon mating (their own and that of the male they mate with). Egg genotypes are then determined by the parental genotypes and inheritance pattern as provided in the genetic inheritance module. The adult equilibrium population size, *N*, in a given habitat patch is used to determine the carrying capacity of that patch for larvae, *K*, which in turn determines the degree of additional density-dependent mortality at the larval stage in that patch. Following Deredec *et al.* [25], this is described by an equation of the form: 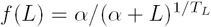, where *L* is the number of larvae in the patch, *T*_*L*_ is the duration of the larval stage, and *α* is a parameter describing the strength of density dependence. Further details on the mathematical formulation of the lumped-age class model and its generalization to an arbitrary number of genotypes using tensor algebra are provided in the S1 File.

**Fig 2.**
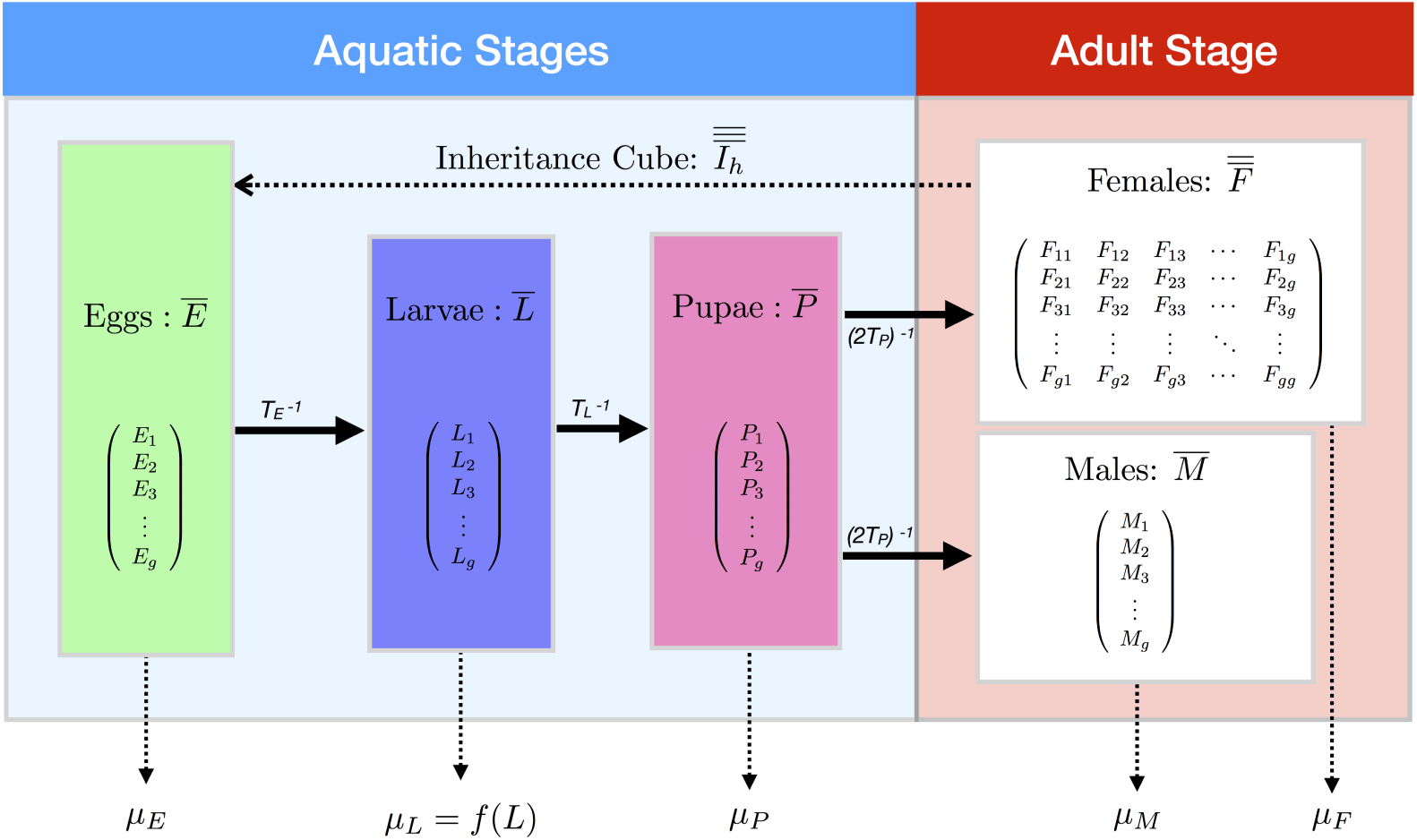
Mosquito life history module. Life history is modeled according to an egg (E)-larva (L)-pupa (P)-adult (M for male, F for female) life cycle in which density dependence occurs at the larval stage and autonomous mobility occurs at the adult stage. Genotypes are tracked across all life stages and are represented by the subscript *i ∈*{1, *…, g*}. E.g. *M*_*i*_ represents the number of adult males having the *i*th genotype, and so on for other life stages and genotypes. Females are modeled as mating once upon emergence and hence obtain a composite genotype - their own and that of the male they mate with. Egg genotypes are determined by the adult female’s composite genotype and the inheritance pattern, which is specific to the gene drive system under consideration.

The MGDrivE framework currently applies to any species having an egg-larva-pupa-adult life history and for which density-dependent regulation occurs at the larval stage. Switching between species can be achieved by altering the parameter values that describe this module when initializing an MGDrivE simulation. The input variables for this module currently include: a) the number of eggs produced per adult female per day, b) the durations of the egg, larval and pupal juvenile life stages, c) the daily mortality risk for the adult life stage, and d) the daily population growth rate (in the absence of density-dependent mortality). The daily density-independent mortality risks for the juvenile stages are assumed to be identical and are chosen for consistency with the daily population growth rate. Default life history parameter values are shown in Table 2 for three species of interest: a) *An. gambiae*, the main African malaria vector, b) *Ae. aegypti*, the main vector of dengue and Zika virus, and c) *Ceratitis capitata*, a worldwide agricultural crop pest. In some cases, life history parameters will be modified in genotype-specific ways by the gene drive construct, and such modifications are efficiently accommodated within this framework via tensor operations. A noteworthy limitation of the current version of the modeling framework is that equilibrium population size remains constant over time, which prevents the user from determining the optimal seasonal timing of a release. This limitation will be addressed in the next released version of MGDrivE.

**Table 2.** Life history module parameter values for three species of interest (at a temperature of 25 Celsius).

| Parameter | Symbol | *Ae. aegypti* | *An. gambiae* | *C. capitata* |
| --- | --- | --- | --- | --- |
| Egg production per female ($\text{day}^{-1}$) | $\beta$ | 20 [36] | 32 [37] | 20 [38] |
| Duration of egg stage (days) | $T_E$ | 5 [39] | 1 [37] | 2 [38] |
| Duration of larval stage (days) | $T_L$ | 6 [39] | 13 [37] | 6 [38] |
| Duration of pupa stage (days) | $T_P$ | 4 [39] | 1 [37] | 10 [38] |
| Daily population growth rate ($\text{day}^{-1}$) | $r_M$ | 1.175 [40] | 1.096 [41] | 1.031 [42] |
| Daily mortality risk of adult stage ($\text{day}^{-1}$) | $\mu_M, \mu_F$ | 0.090 [43–45] | 0.123 [41] | 0.100 [46] |

### 3. Landscape

The landscape module describes the distribution of mosquito metapopulations in space, with movement through the resulting network determined by dispersal kernels. Metapopulations are randomly mixing populations for which the equations of the lumped age-class model apply. The resolution of the metapopulations (in terms of size) should be chosen according to the dispersal properties of the insect species of interest and the research question being investigated. *Ae. aegypti* mosquitoes, for instance, are thought to be relatively local dispersers, often remaining in the same household for the duration of their lifespan [47]. For modeling the fine-scale spread of gene drive systems in this species, metapopulations the size of households may be appropriate. *An. gambiae* mosquitoes, on the other hand, are thought to display moderate dispersal on the village scale and infrequent inter-village movement [48]. This would suggest villages as an appropriate metapopulation unit; however other levels of aggregation are also possible, in both cases, depending on the level of resolution required from the simulations and the computational power available to the user.

Once the metapopulation size has been decided upon and the metapopulations have been enumerated, MGDrivE accepts a list of coordinates and equilibrium adult population sizes associated with each. In the resulting network structure, nodes represent randomly-mixing metapopulations and edges represent movement of mosquitoes from one metapopulation to any other in the network (Figure 3). Movement between metapopulations is limited to the adult life stage. By default, movement rates between metapopulations are derived from a zero-inflated exponential dispersal kernel, the degree of zero-inflation and mean dispersal distance of which may be defined by the user. That said; the movement kernel may be expanded arbitrarily to account for barriers to movement such as roads [47] and other factors without altering the overarching model structure. Movement rates between nodes are then used to calculate a matrix of node transition probabilities, which is incorporated in the tensor algebraic model formulation described in the S1 File.

**Fig 3.**
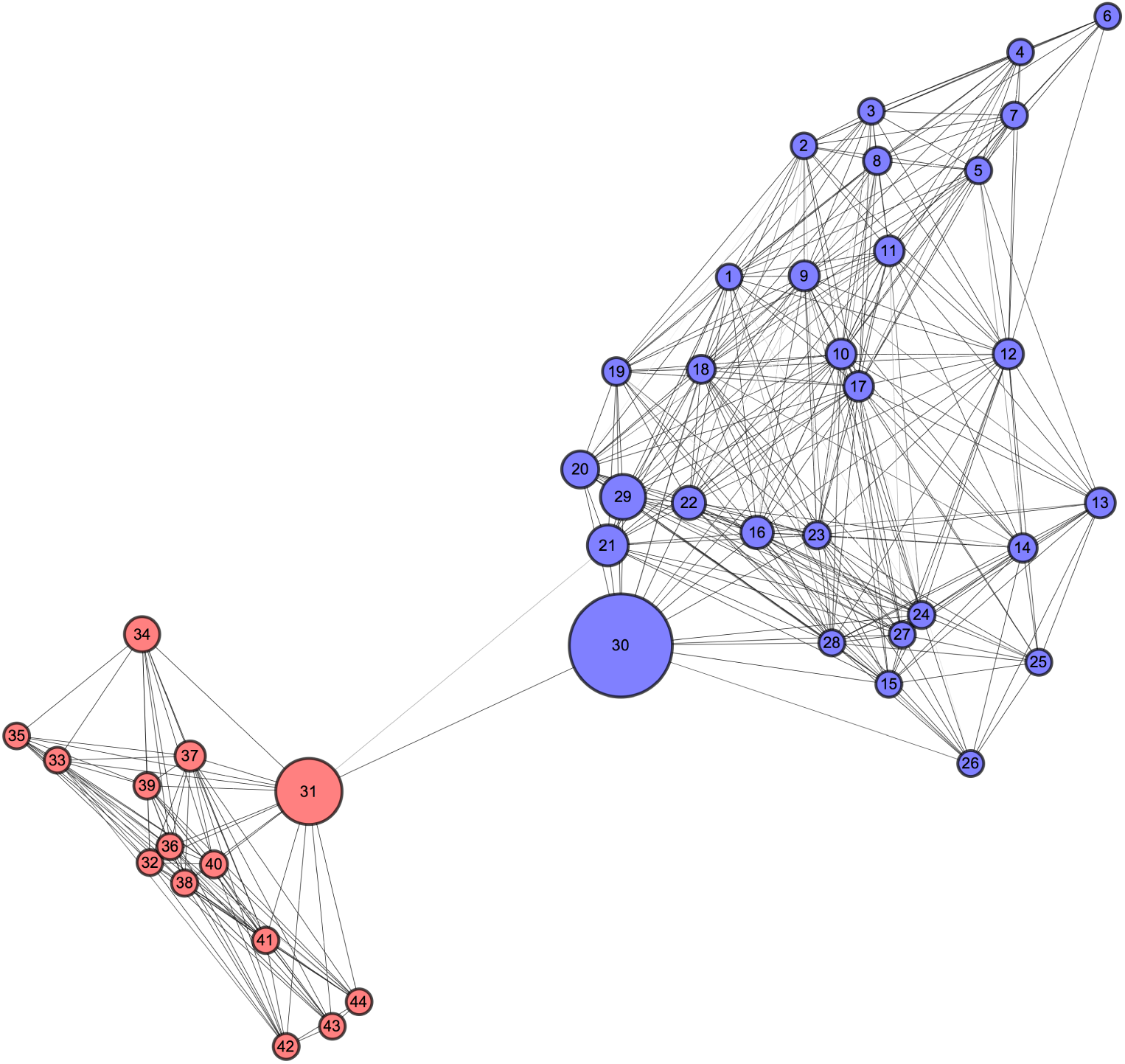
Landscape module. Insects are distributed as metapopulations, here depicted by nodes, each having their own coordinates and population size. Movement between metapopulations is derived from a defined dispersal kernel and is depicted here by edges between nodes. The example “tale of two cities” scenario allows both spread within and between communities to be explored. Here, nodes are colored according to their community (as detected by the DBSCAN clustering algorithm [49]), with sizes proportional to their “betweenness centrality” - a measure of their connectedness to other nodes in terms of number of shortest paths that flow through them [50].

Finally, with the inheritance, life history and landscape modules in place, any type of release can be simulated by increasing the number of insects having the released sex and genotype at a specific metapopulation and time. As demonstrated in the following software use example, input variables are provided for: a) release size, b) number of releases, c) time of first release, d) time between releases, e) metapopulation of release, and f) sex and genotype of released insects.

## Deterministic vs. Stochastic Simulations

Simulations in MGDrivE can be run either in deterministic or stochastic form. Deterministic simulations are faster and less computationally intensive; however, stochastic simulations capture the probabilistic nature of chance events that occur at low population sizes and genotype frequencies. For instance, a stochastic model is required to understand the chance of population elimination following releases of insects carrying a population-suppressing homing system in the context of rarely generated resistant alleles [26]. In the stochastic implementation of MGDrivE, daily egg production follows a Poisson distribution, offspring genotype follows a multinomial distribution informed by parental genotypes and the inheritance pattern of the gene drive system, mate choice follows a multinomial distribution determined by adult genotype frequencies, and survival and death events follow binomial distributions at the population level. When interpreting stochastic models, many simulations should be run to understand the range of outputs possible for a given model realization.

## Two Example MGDrivE Simulations

To demonstrate how the MGDrivE framework can be used to initialize and run a simulation of a gene drive system through a network of connected metapopulations, we describe the application of the package to two CRISPR/Cas9-based homing gene drive strategies: a) driving a disease-refractory gene into a population [7], and b) disrupting a gene required for female fertility and hence suppressing a population [30]. In both cases, we consider a population of *Ae. aegypti* mosquitoes having the bionomic parameters provided in Table 2 and distributed through the network landscape depicted in Figure 3. To demonstrate the functionality of the MGDrivE package, we model the population replacement strategy (i.e. replacing the population with a disease-refractory one) using the deterministic implementation, and model the population suppression strategy using the stochastic implementation. The stochastic implementation is more relevant to population suppression as it can capture rare resistant allele generation and the possibility of population extinction. In both cases, we include the generation of in-frame and out-of-frame or otherwise costly resistant alleles [28, 51] and parameterize the gene drive model based on recently engineered constructs [7, 30].

### 1. Population Replacement

We begin by modeling a CRISPR/Cas9-based homing construct similar to that engineered by Gantz et al. [7]. This was the first CRISPR-based homing construct demonstrated in a mosquito disease vector – namely, *Anopheles stephensi*, the main urban malaria vector in India. For this construct, homing and resistant allele generation were shown to occur at different rates in males and females, and there were large fitness reductions associated with having the homing construct. We consider a homing efficiency of 90% in males and 50% in females – i.e. 90% of wild-type (h) alleles are converted to homing (H) alleles in the germline of Hh males, and 50% of h alleles are converted to H alleles in the germline of Hh females. A third of the remaining h alleles in Hh individuals are converted to in-frame resistant alleles (R), and the remainder are converted to out-of-frame or otherwise costly resistant alleles (B) due to error-prone copying during the homing process [51]. Female fecundity and male mating fitness are reduced by 25% per H or R allele and by 50% per B allele.

The code for this simulation (Code samples 1-3) can be entered directly in R, and the details of the various functions used are described in the online documentation available at https://marshalllab.github.io/MGDrivE/docs/reference/. The general workflow for the simulation is shown in Figure 4. We begin by loading the MGDrivE package in R and choosing the working and output directories. The output directory should be a dedicated directory for MGDrivE simulation output, to avoid interfering with other files. We then choose between the deterministic and stochastic implementation of the model framework – in this case the deterministic version. Next, we specify the bionomic parameters of the species we are modeling – in this case, *Ae. aegypti*, whose default life history parameters are provided in Table 2. Following this, we define the landscape through which we will model the spread of the drive system. We begin by loading a CSV file containing the coordinates (longitude and latitude) of the metapopulations in Figure 3. A function is then applied that computes daily movement rates between each of the metapopulations based on a zero-inflated exponential dispersal kernel, the parameters for which we provide. Equilibrium adult population sizes can be provided for each of the metapopulations; however in this case, we assume these are identical across all metapopulations and provide a single population size (Code sample 1).

**Fig 4.**
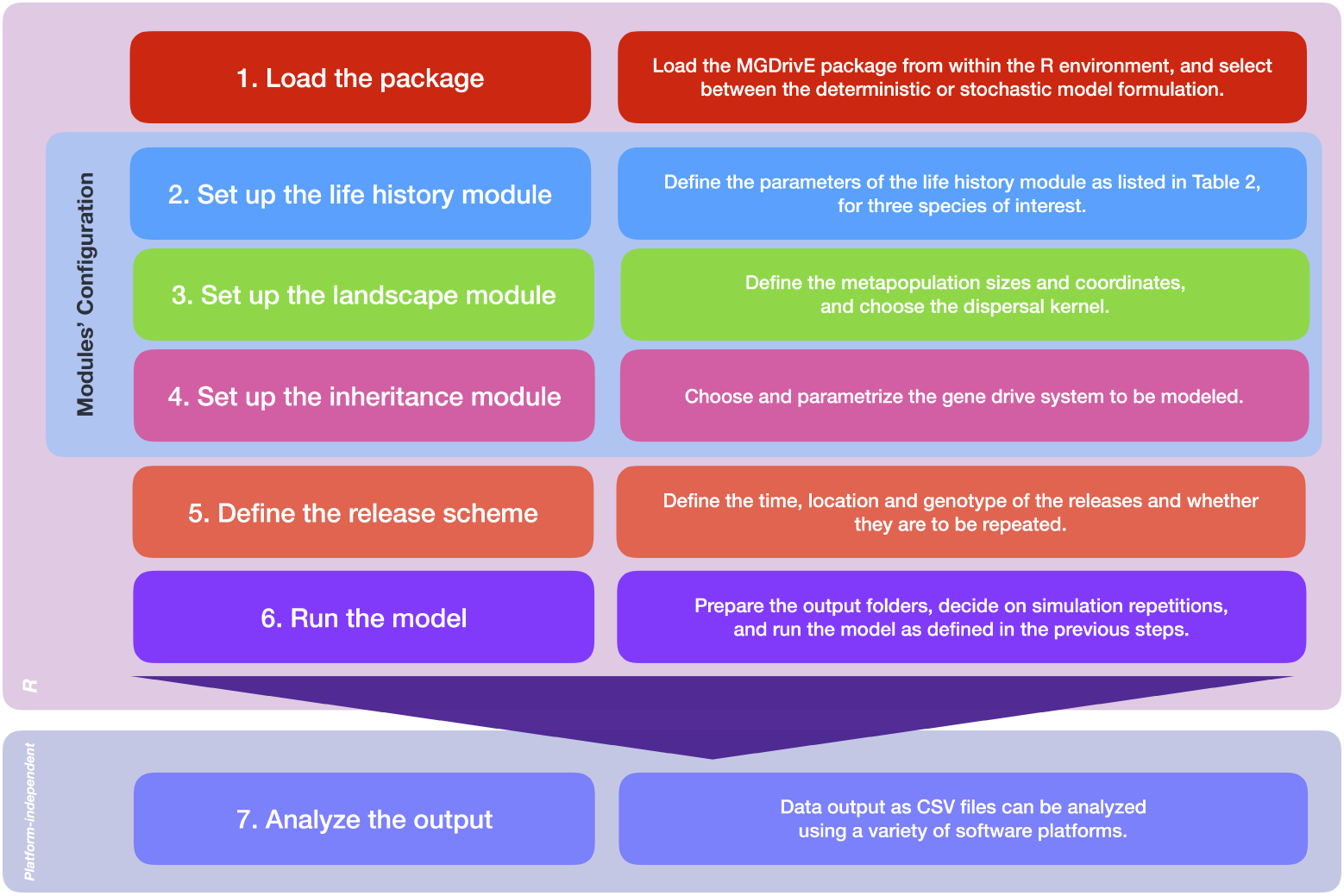
Workflow of an MGDrivE simulation.

**Code sample 1.**
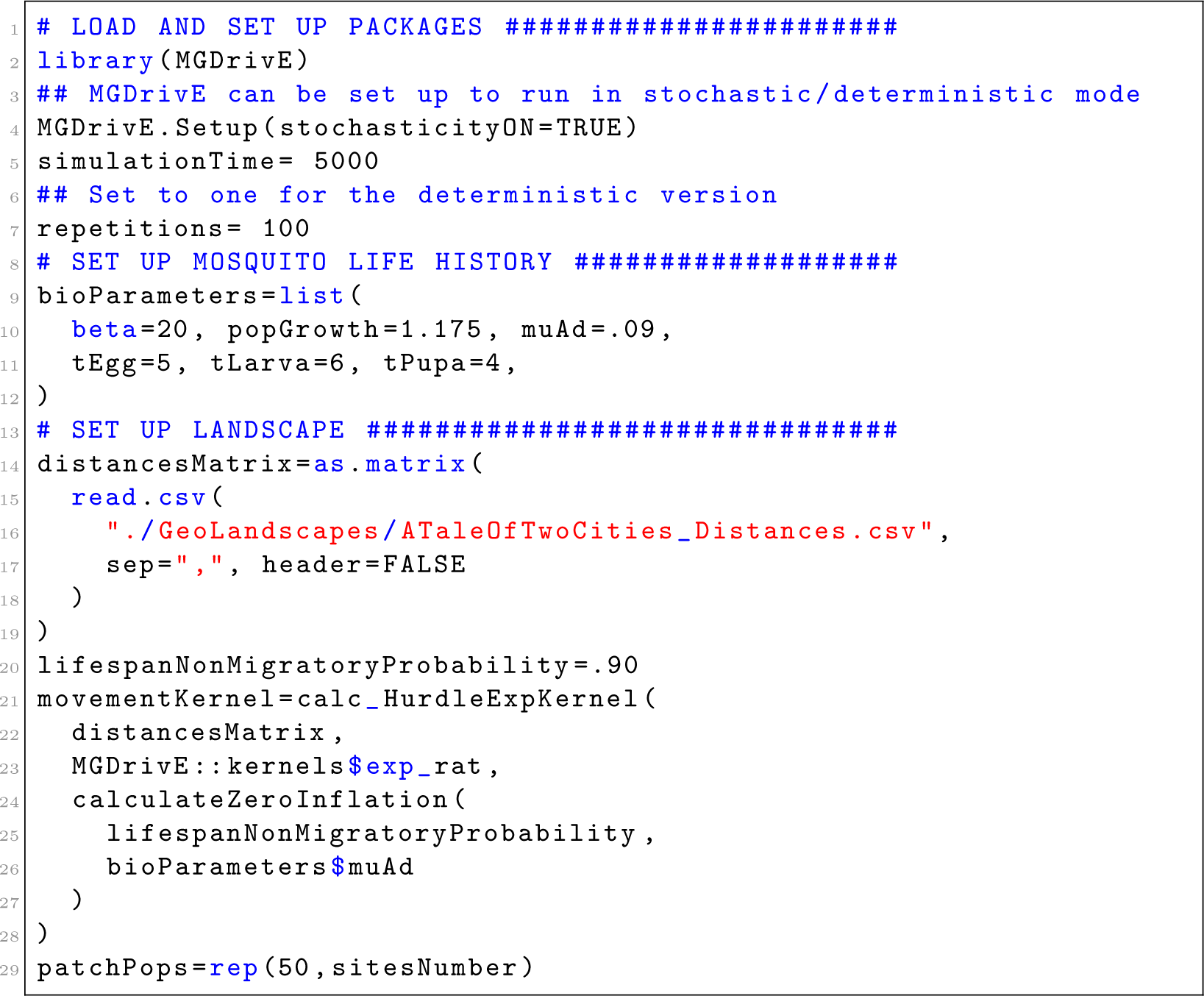
Loading the package and setting up the life history and landscape modules.

With our life history and landscape modules defined and parameterized, we now specify the gene drive system and release strategy we intend to model (Code sample 2). We use a pre-specified inheritance cube function, “Cube HomingDrive()”, that models the inheritance pattern of a homing-based gene drive system. The input options for this function can be seen by typing “?Cube HomingDrive()” at the command prompt. We specify the sex-specific homing rates, resistant allele generation rates, and genotype-specific fitness effects as described earlier based on the construct engineered by Gantz et al. [7]. We then specify the release scheme by generating a list containing: a) the release size, b) number of releases, c) time of first release, and d) time between releases. This is then incorporated into a vector also specifying the inheritance cube and the sex and genotype of the released insects. Finally, the metapopulations in which the release takes place are specified. With the simulation framework now fully specified, the model is now ready to run (Code sample 3).

**Code sample 2.**
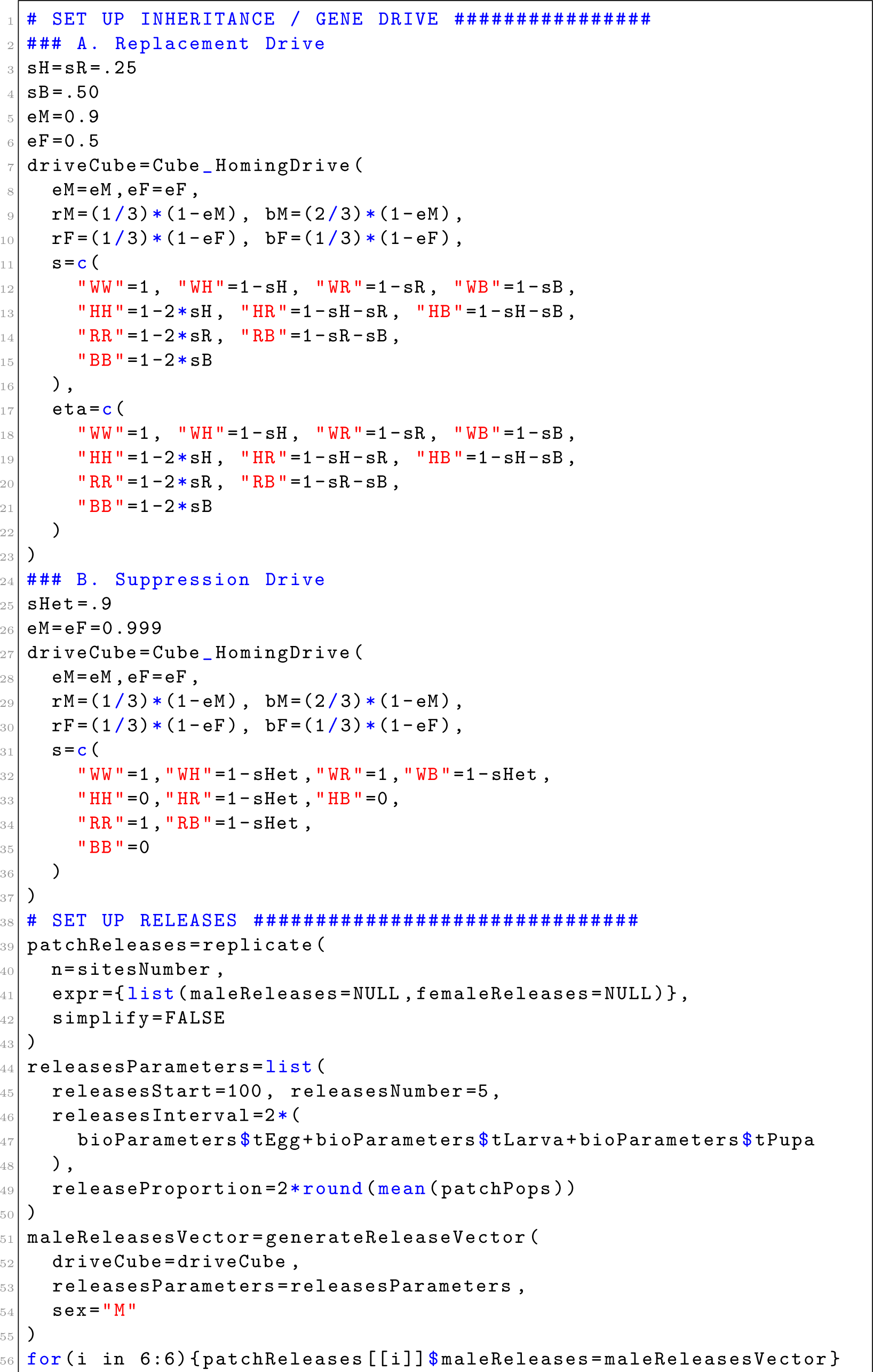
Setting up the inheritance/gene drive module and defining the release scheme. Here, code is shown for both: A) homing-based replacement drive, and B) suppression drive. Only one of these should be selected when running the simulation.

**Code sample 3.**
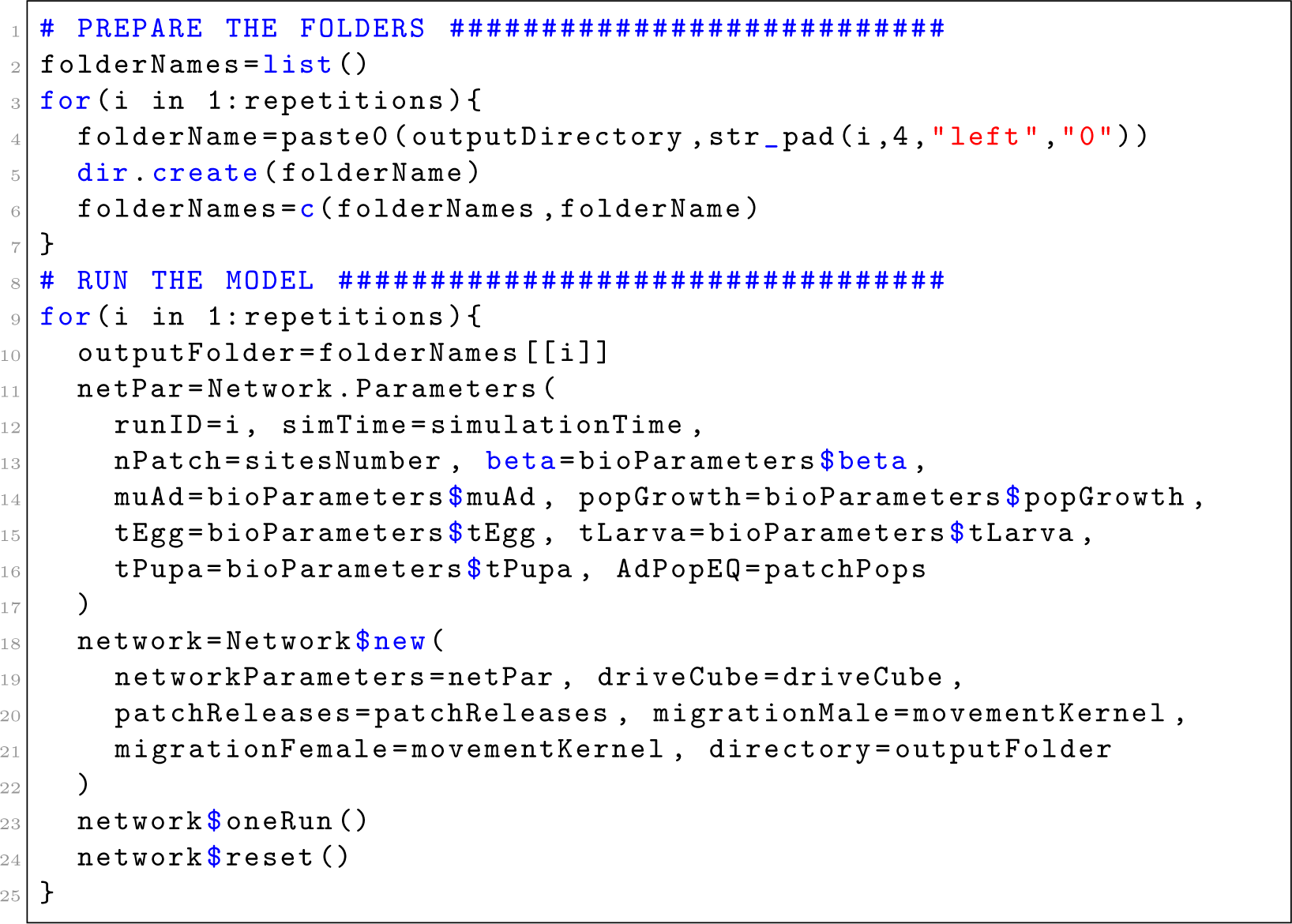
Preparing output folders and running the model. It is recommended to store simulation files for each run in its own separate folder.

### 2. Population Suppression

As a second example, we demonstrate the application of the MGDrivE package to model a population suppression homing construct similar to that engineered by Hammond et al. [30]. For this construct, the homing system targets a gene required for female fertility, causing females lacking the gene (those having the genotypes HH, HB and BB) to be infertile, and inducing a large fecundity reduction of 90% in females only having one functioning copy of the gene (those having the genotypes Hh, HR, hB and RB). The homing efficiency is very high – 99.9% in both males and females – with a third of the remaining h alleles in Hh individuals being converted R alleles and the remainder being converted to B alleles. This is similar to the first CRISPR-based homing construct demonstrated in *An. gambiae*, although with a higher homing efficiency that could be achieved through guide RNA multiplexing [26]. Lines of code that differ for this system are shown in Code sample 2. We choose the stochastic implementation of the model framework this time, and while the same inheritance cube function applies, it’s parameters differ – namely, homing and resistant allele generation rates, and genotype-specific fitness effects.

### Output Analysis

In the current version of MGDrivE, simulation results are output as CSV files, which enables the user to analyze results in any platform of their choice – R, Python, Mathematica, etc. What the user decides to plot will depend on the number of possible genotypes, whether the male-to-female ratio is altered, whether the population is suppressed, and the spatial structure of the landscape through which drive occurs. If the number of genotypes is large, for instance, then allele abundance may provide a more manageable output than that of genotypes.

In Figure 5, we display a potential visualization scheme produced in Mathematica for the population replacement and suppression simulations described above (additionally, videos for both simulations running in the spatial networks can be accessed in the supplementary information: S1 Video and S2 Video). As there are four alleles for both systems (the homing allele, H, the wild-type allele, h, and the two resistant alleles, R and B), we depict their abundance in the Figures 5A and 5B and their frequency in Figures 5C and 5D, with time on the horizontal axis and metapopulation number on the vertical axis. For population replacement (Figures 5A and 5C), we see the gene drive system (H) spread through the population, and the in-frame resistant allele (R) accumulate to a small extent. This occurs because the R allele has neither a fitness cost nor benefit relative to the H allele once it has saturated the population, while the B allele is selected against due to its inherent selective disadvantage. For population suppression (Figures 5B and 5D), we see the gene drive system (H) spread through the population at the same time as it induces suppression due to its impact on female fertility. Eventually, we see an in-frame resistant allele (R) emerge and spread into the population due to its selective advantage over both the wild-type and homing alleles. Also visible in Figure 5 is the slightly extended time it takes for both homing systems to spread through the second population cluster visible in the metapopulation landscape depicted in Figure 3.

**Fig 5.**
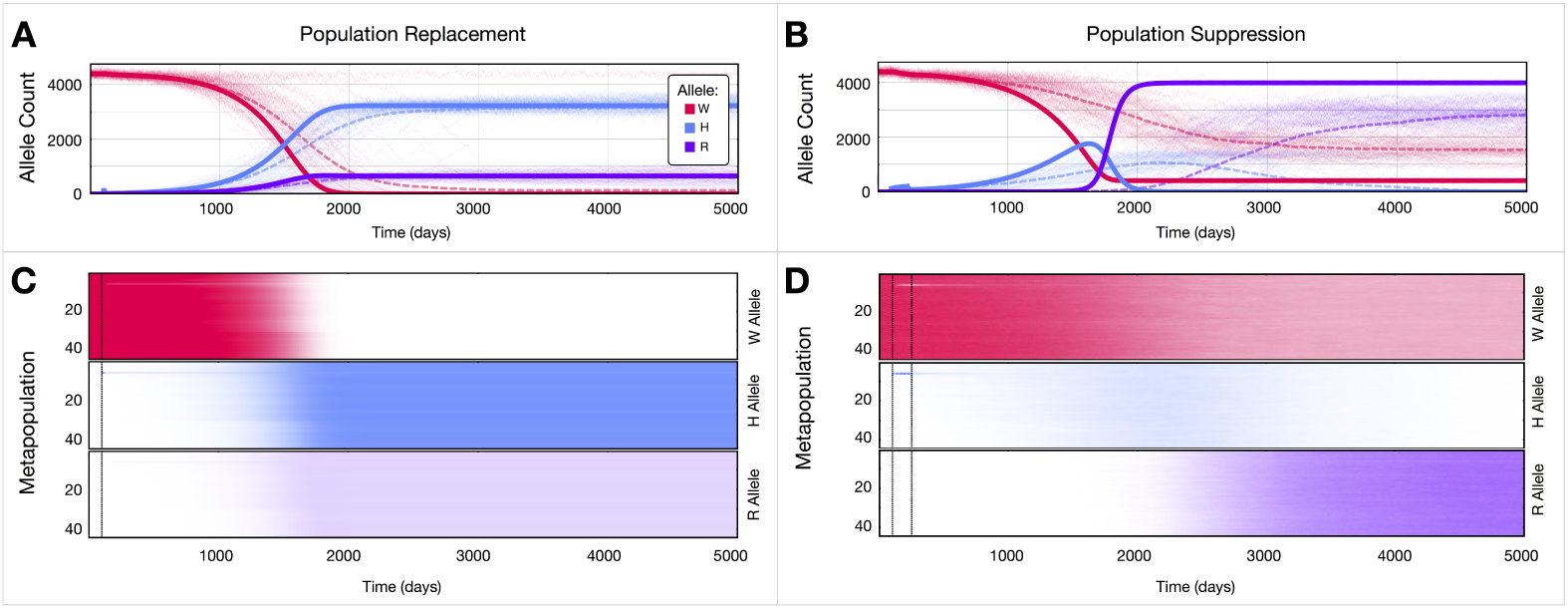
Example MGDrivE simulations for CRISPR-based homing constructs. In both cases, an *Aedes aegypti* population is simulated having the bionomic parameters in Table 2 and distributed through the landscape depicted in Figure 3. Deterministic simulations are denoted by solid lines in panels A and B, while stochastic simulations are denoted by thin lines, each corresponding to the output of a single simulation, and dotted lines, corresponding to the mean of 100 stochastic simulations. **A.** A population replacement homing construct that drives a disease-refractory gene into the population is simulated having a homing efficiency of 90% in males and 50% in females. Wild-type (h) alleles that are not converted to homing (H) alleles in the germline of Hh heterozygotes are cleaved and converted to either in-frame (R) or out-of-frame (B) resistant alleles. Female fecundity and male mating fitness are reduced by 25% per H or R allele and by 50% per B allele. A single release of 100 HH females at node 6 is modeled. As the homing allele (blue) is driven into the population, the wild-type allele (red) is eliminated, and the in-frame resistant allele (purple) accumulates to a population frequency of 17%. Stochasticity slightly slows the allele frequency trajectories, on average, and introduces variability around the mean output. **B.** A population suppression homing construct that interferes with a gene required for female fertility is simulated having a homing efficiency of 99.9% in both females and males. Wild-type alleles that are not converted to homing alleles in the germline of Hh heterozygotes are cleaved and converted to either in-frame or out-of-frame resistant alleles. Females without a copy of the h or R allele are infertile, while females having only one copy of the h or R allele have a 90% fecundity reduction. Five releases of 100 HH females at node 6 are modeled. As the homing allele (blue) is driven into the population, it suppresses the population due to its impact on female fertility. Eventually, an in-frame resistant allele (purple) emerges and leads the population to rebound due to its selective advantage over both wild-type and homing alleles. In the deterministic model output, the in-frame resistant allele spreads to fixation; however in the stochastic model output, the homing allele is sometimes lost from the population and, as a result, the selective advantage of the in-frame resistant allele is lost, causing it to equilibriate at a lower population frequency. Stochasticity also significantly slows the mean allele frequency trajectories, as well as introducing variability around the mean. **C-D.** Here, population frequencies of the wild-type, homing and in-frame resistant alleles are shown in each metapopulation over time for a deterministic model of the population replacement construct (panel C) and a stochastic simulation of the population suppression construct (panel D). Out-of-frame resistant alleles are omitted due to their low frequencies in both simulations. Dashed vertical lines represent the beginning and end of the releases.

## Availability and Future Directions

As of the date of publication, we are releasing MGDrivE version 1.0 (“Rise and Shine”), available at our permanent github repository at: https://github.com/MarshallLab/MGDrivE. The source code is available under the GPL3 License and is free for other groups to modify and extend as needed. Mathematical details of the model formulation are available in the S1 File, and documentation of all MGDrivE functions are available at the project’s github repository at https://marshalllab.github.io/MGDrivE/docs/reference/. To run the software, we recommend using R version 3.4.4 or higher.

We are continuing development of the MGDrivE software package, and welcome suggestions and requests from the research community regarding future directions. The field of gene drive has been moving extremely quickly, especially since the discovery of CRISPR-based gene editing, and we intend the MGDrivE package to provide a flexible tool capable of modeling novel inheritance-modifying constructs as they are proposed and become available. Future functionality that we intend to incorporate into the software includes: a) “shadow drive”, in which the Cas9 enzyme is passed on to the offspring even if the gene expressing it is not [51], b) life history models encompassing a more diverse range of insect disease vectors and agricultural pests, and c) populations that vary in size seasonally or in response to environmental drivers such as temperature and rainfall. The incorporation of environmental drivers will allow both seasonal trends and short-term fluctuations to be accommodated within the same framework.

Alongside our population-based model, we are also developing a corresponding individual-based model that is capable of modeling multi-locus systems for which the number of possible genotypes exceeds the number of individuals in the population. This will enable us to efficiently model confineable systems such as daisy-drive involving several loci [27], and multiplexing schemes in which a single gene is targeted at multiple locations with separate guide RNAs to reduce the rate of resistant allele formation [52].

## Supporting information

**S1 Video. Population replacement use example.** A visualization of the homing-based population replacement simulation.

**S2 Video. Population suppression use example.** A visualization of the homing-based population suppression simulation.

**S1 Text. Introduction to the modeling framework.** A description of the mathematical equations that govern the inheritance, life history and landscape modules.

## Acknowledgments

The authors would like to thanks Drs. Omar Akbari, Ethan Bier and Anthony James for discussions on gene drive architectures and molecular biological considerations, and Drs. Gregory Lanzaro, Yoosook Lee and Gordana Rašić and Ms. Partow Imani for discussions on mosquito ecology, life history and dispersal behavior. This work was supported by a DARPA Safe Genes Program Grant (HR0011-17-2-0047) awarded to JMM and funds from the UC Irvine Malaria Initiative and Innovative Genomics Institute awarded to JMM.

## MGDrivE: Mosquito Gene Drive Explorer

## 1 Introduction

The advent of CRISPR/Cas9-based gene editing technology and its application to the engineering of gene drive systems has led to renewed excitement in the use of genetics-based strategies to control mosquito vectors of human diseases and insect agricultural pests. Applications to control mosquito-borne diseases have gained the most attention due to the major global health burden they pose through much of the world [1,2] and the difficulty of controlling them using currently-available tools [3]. The recent engineering of a gene drive system in *Drosophila suzukii* [4], a major insect agricultural crop pest, and the difficulty of controlling insect crop pests using existing tools, has led to growing enthusiasm for the application of these tools to other insect species too.

The versatility of this technology has also enabled a wide range of gene drive architectures to be realized [5]. Prior to the advent of CRISPR, homing endonuclease genes (HEGs) were envisioned to cleave a specific target site lacking the HEG and to be copied to this site by serving as a template for homology-directed repair, effectively converting a heterozygote into a homozygote and biasing inheritance in favor of the HEG [6]. A vast range of additional approaches for biasing inheritance are now being proposed, including several threshold-dependent systems that may permit confineable and reversible releases [7], and remediation systems that could be used to remove effector genes and possibly entire drive systems from the environment in the event of unwanted consequences [8].

Understanding how these systems are expected to behave in real ecosystems requires a flexible modeling framework that can accommodate a range of inheritance patterns, specific details of the species into which the constructs are to be introduced, and details of the landscape through which spatial spread would occur. To this end, we present **MGDrivE** (Mosquito Gene Drive Explorer): a flexible simulation framework designed to investigate the population dynamics of a variety of gene drive systems and their spread through spatially-explicit populations of mosquito species and other insect species.

A key strength of the **MGDrivE** framework is its modularity [Figure 1]. A genetic inheritance module allows the inheritance dynamics of a wide variety of drive systems to be accommodated. An independent population dynamic module allows the life history of a variety of mosquito disease vectors and insect agricultural pests to be accommodated. Thirdly, a landscape module accommodates the distribution of insect metapopulations in space, with movement through the resulting network determined by dispersal kernels. The model can be run in either a deterministic or stochastic form, allowing the chance events that occur at low population or genotype frequencies to be simulated.

**Figure 1:**
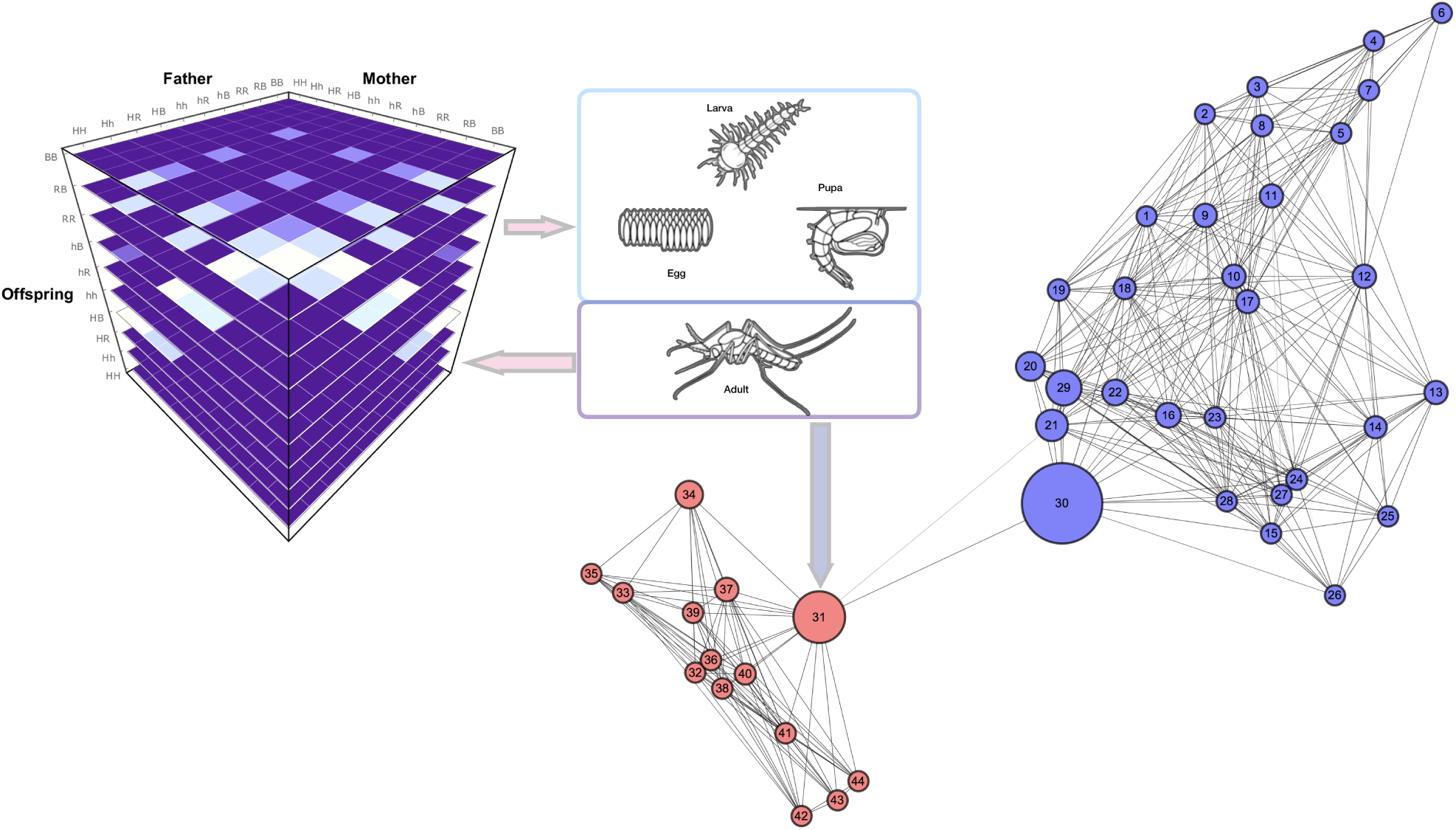
Interaction of the inheritance, life history and landscape components of the **MGDrivE** modeling framework. The inheritance module (left) informs the distribution of egg genotypes given two parent genotypes. The life history module describes the population dynamics of the mosquito life stages (top, center). The landscape module (right & lower center) describes movement of mosquitoes between metapopulations in which the population dynamic equations apply.

What separates **MGDrivE** from other models is its basis in a parsimonious set of mathematical equations that accommodate these three modules and can be efficiently modified to accommodate a wide array of genetics-based systems, insect species and landscapes of interest. This is achieved by treating the population dynamics within a variable-dimension tensor algebraic framework. As different genetics-based systems are modeled, the dimensionality of the equations changes; but the population dynamic equations remain the same. In the following sections, we describe this tensor-modeling framework in more detail.

## 2 Notation

We begin by defining some of the notation conventions we will follow for the written description of the model.

- As our framework is based on tensor operations, we will use overlines to denote tensor dimension.
- As our framework is based on discrete-time difference equations, we will use subscript square brackets to indicate time (measured in days). E.g., 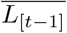 denotes the genotype-specific larval population size at time (or day) *t* − 1.
- Matrices follow a “row-first” indexing order. I.e. row *i*, column *j*.
- Hadamard products (∘) denote operations for which entries are multiplied entry-by-entry. These operations apply to tensors, matrices and vectors that have the same dimensionality.
- Cross products (*×*) represent standard matrix/vector multiplication.
- Outer products (*⊗*) between two vectors, a and 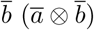, produce matrices, 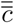, whose entries are given by *c*_*ij*_ = *a*_*i*_*· b*_*j*_.
- We use a dot (*·*) to denote scalar multiplication.

## 3 Inheritance module

### 3.1 Inheritance cubes

The fundamental module for modeling gene drive dynamics is that describing genetic inheritance. In **MG-DrivE**, this is embodied by a three-dimensional tensor, 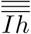, referred to as an “inheritance cube”. Each gene drive system has a unique R file containing the three-dimensional inheritance cube. The first and second dimensions of the inheritance cube refer to the maternal and paternal genotypes, respectively, and the third dimension refers to the offspring genotype. The cube entries for each combination of parental and offspring genotypes represent, for each parental pairing, the proportion of offspring that are expected to have each genotype. For a given set of parents, these entries should sum to one, as fitness and viability are accommodated separately. Handling this within a tensor framework allows an arbitrary number of genotypes to be efficiently accommodated while maintaining the same population dynamic equations.

### 3.2 Maternal and paternal genotypes

Before describing how inheritance is implemented within the **MGDrivE** framework, we first describe how genotype information is accommodated for female and male adults. Adult females and males are treated differently in this framework, since it is assumed that female mosquitoes only mate once, while male mosquitoes may mate throughout their lifetime. Males therefore have their own genotype, indexed in vector form by *i*, while females have a composite genotype consisting of their own genotype, *i*, and the genotype of the male with whom they mated, *j*.

For a genetic system consisting of *g* genotypes, the number of adult males having each genotype is denoted by the vector, 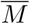, the *i*th entry of which (*M*_*i*_) denotes the number of adult males having the *i*th genotype. I.e.:

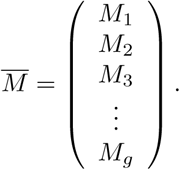

In the population dynamic framework, this genotype indexing notation is consistent throughout.

As adult females have a composite genotype, the number of adult females having each composite genotype is denoted by the matrix, 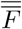, the *i*th row of which denotes the adult female’s own genotype, and the *j*th column of which denotes the male with whom they mated. The entry in the *i*th row and *j*th column of this matrix (*F*_*ij*_) denotes the number of adult females having the *i*th genotype and having mated with an adult male having the *j*th genotype, i.e.:

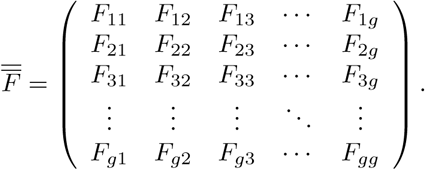

### 3.3 Calculating the expected distribution of offspring genotypes

In the population dynamic framework, the inheritance cube, 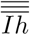, is encountered when eggs are laid by mated females. Included in the calculation of the number of eggs laid per day having each genotype are:

- the number of adult females having each mated genotype, 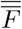,
- the fecundity of adult females having each genotype, 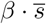, where *β* represents the number of eggs produced per day per wild-type female, and *s* is a vector of genotype-specific multipliers for each female’s own genotype,
- the “inheritance cube” tensor, 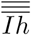, as described earlier, and
- a “viability mask” tensor, 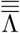, which removes eggs from the pool that are unviable due to the specific combination of maternal, paternal and offspring genotypes.

This latter feature is particularly relevant for toxin-antidote-based gene drive systems such as *Medea* [9], for which wild-type offspring of heterozygous mothers are unviable as they don’t have the antidote to the toxin deposited in the embryo by the mother.

The number of eggs having the *i*th genotype that are oviposited at time *t, E*_*i,*[*t*]_, is then given by:

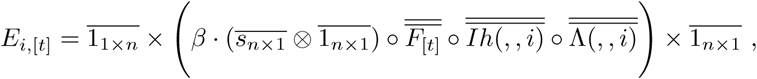

where 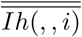 and 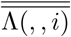 represent the slices of the inheritance cube, 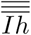, and viability mask tensor, 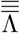, corresponding to the offspring genotype, *i*. This calculation may be repeated for all offspring genotypes, *i ∈*{1, *…, g*}, to produce the corresponding vector, 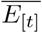. This equation demonstrates the flexibility afforded by the tensor framework, as the equation strucure can be maintained while the dimensionality is changed according to the needs of the inheritance-biasing system.

### 3.4 Currently-available inheritance cubes

In version 1.0 of **MGDrivE**, the following inheritance cubes are provided:

- `Cube Mendelian:` Standard Mendelian inheritance
- `Cube HomingDrive:` Homing-based drive with in-frame and out-of-frame/costly resistance alleles
- `Cube Homing1RA:` Homing-based drive with one type of resistance allele
- `Cube ImmunizingReversalMF:` Homing-based immunizing reversal drive
- `Cube MEDEA:` *Medea* (Maternal effect Dominant Embryonic Arrest)
- `Cube oneLocusTA:` Single-locus version of UD^MEL^
- `Cube twoLocusTA:` Two-locus version of UD^MEL^
- `Cube KillerRescue:` Killer-rescue system for transient gene drive
- `Cube ReciprocalTranslocations:` Reciprocal chromosomal translocations
- `Cube Wolbachia:` *Wolbachia*
- `Cube RIDL:` RIDL (Release of Insects carrying a Dominant Lethal gene)

## 4 Life history module

### 4.1 Lumped age-class model

The equations for the life history module of **MGDrivE** follow from the lumped age-class model of Hancock and Godfray [10] adapted by Deredec *et al.* [11], and later by Marshall *et al.* [5] to describe the spread of a homing-based drive system with resistance alleles through a density-dependent population. In this model, a daily time step is used, and the mosquito life cycle is divided into four life stages - egg, larva, pupa, and adult (both female and male) - denoted by the subscripts “E”, “L”, “P”, “M” and “F”, respectively.

The daily, density-independent mortality rates for the juvenile stages are assumed to be identical (*μ*_*E*_ = *μ*_*L*_ = *μ*_*P*_) and are chosen for consistency with the population growth rate in the absence of density-dependent mortality, *R*_*M*_. I.e.:

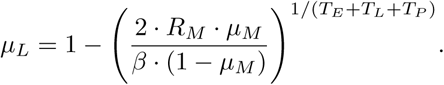

Here, *μ*_*M*_ denotes the mortality rate of adult male (and female) mosquitoes, and *T*_*E*_, *T*_*L*_ and *T*_*P*_ denote the duration of the egg, larval and pupal life stages. The probability of surviving any of the juvenile stages in a density-independent setting, *θ*_*x*_, is given by:

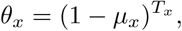

where *x ∈*{*E, L, P*}; however additional density-dependent mortality, 1 − *f* (*L*), occurs at the larval stage. We use a density-dependent equation of the following form to model this:

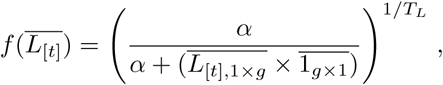

where *α* is a parameter influencing the strength of density-dependence. The *α* parameter is chosen to produce the desired equilibrium density of adult mosquitoes in the population, *N*_*eq*_. I.e.:

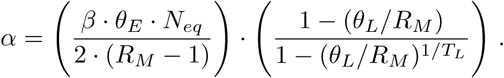

Of note, the daily population growth rate, *r*_*M*_, is related to the population growth rate per generation, *R*_*M*_, according to:

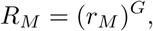

where *G* is the mosquito generation time, and is given by:

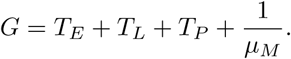

### 4.2 Population dynamics of each life stage

With this framework in place, the dynamics of the population can be described by equations for the number of larvae and adults belonging to each genotype at time *t*. The number of larvae at time *t* is needed to determine the strength of density-dependence and, partitioned by genotype, is given by:

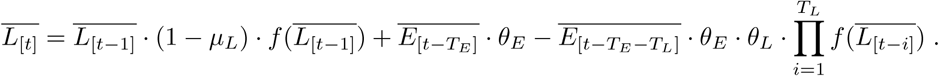

Here, the first term accounts for survival of larvae (denoted at time *t* by 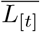) from one day to the next, the second term accounts for newly hatching eggs having each genotype, and the third term accounts for development of larvae into pupae for larvae having each genotype.

Adult females and males are treated slightly differently in this framework since it is assumed that female mosquitoes only mate once, soon after emergence, while male mosquitoes may mate throughout their lifetime. The number of adult males having each genotype at time *t* is given by:

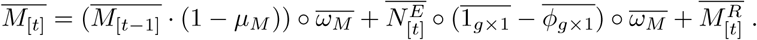

Here, the first term accounts for survival of adult males (denoted at time *t* by 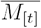) from one day to the next, the second term accounts for development of pupae into adult males, and the third term accounts for males released into the population at time *t*. Some new terms have been introduced here:

- 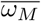 is a genotype-specific multiplier on the daily survival probability for adult males (1 corresponds to default survival/no additional mortality, 0.5 corresponds to a 50% reduction in daily survival)
- 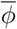 represents the proportion of emerging adult mosquitoes that are female (0.5 corresponds to equal numbers of females and males, where as 0.75 corresponds to a ratio of 3 emerging females to 1 emerging male)

Additionally, 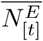 represents the number of emerging adults (female and male) at time *t*, neglecting the genotype-specific multiplier on adult survival on the day or emergence:

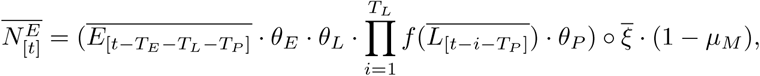

where 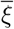 is a genotype-specific multiplier on pupal survival representing pupatory success (1 corresponds to default pupatory success, 0.5 corresponds to a 50% reduction in pupatory success).

Females, on the other hand, are assumed to mate only once, and on the same day they emerge. As far as offspring are concerned, they therefore have attributes for both their own genotype and the genotype of the male with whom they mated. The number of adult females having each genotype at time *t* is given by:

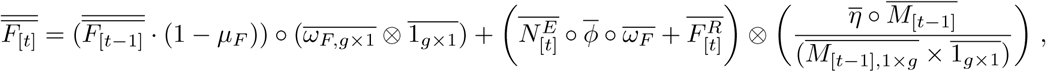

Here, the first term accounts for survival of adult females (denoted at time *t* by 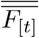) from one day to the next, the first term in the first set of large brackets accounts for development of pupae into adult females, the second term in the first set of large brackets accounts for adult females released into the population at time *t*, and the term in the second set of large brackets accounts for mating of newly emerging and released adult females with adult males having each genotype. Wild-type adult female mortality is set equal to wild-type adult male mortality, i.e. *μ*_*F*_ = *μ*_*M*_.

A new term has been introduced here, 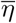, which represents the mating competitiveness of each male genotype, relative to wild-type (1 corresponds to equal mating competitiveness as compared to a wild-type male, while 0.5 corresponds to half the mating competitiveness of a wild-type male). A term corresponding to relative fecundity of females having each genotype relative to wild-type females, 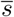, is described in the “Inheritance module” in the equation for 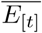 (the number of eggs having each genotype that are oviposited at time *t*).

### 4.3 Releases of adult females and males

Releases of adult females and males are accommodated within this framework through the 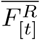 and 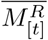 vectors, which denote the number of adult females and males, having each genotype, released into the population at time *t*. The **MGDrivE** software package includes flexible functionality to allow releases at regular or irregular intervals and with different genetic compositions (so long as the released genotypes are included in the inheritance cube).

## 5 Landscape module

The landscape module describes the distribution of mosquito metapopulations in space, with movement through the resulting network determined by dispersal kernels. Metapopulations are randomly mixing populations for which the equations of the lumped age-class model apply, the size of which should be chosen according to the dispersal properties of the insect species of interest and the research question being investigated. Once this has been decided upon, **MGDrivE** accepts a list of coordinates and equilibrium adult population sizes associated with each.

In the resulting network structure, nodes represent randomly-mixing metapopulations and edges represent movement of mosquitoes from one metapopulation to any other in the network. Movement between metapopulations is limited to the adult life stage. By default, movement rates between metapopulations are derived from a zero-inflated exponential dispersal kernel such that, for metapopulations *i* and *j* a distance *d*_*ij*_ apart, the rate of movement between the metapopulations is:

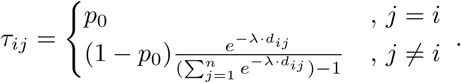

Here, *n* represents the number of metapopulations in the landscape, *p*_0_ represents the probability that a mosquito remains in the same metapopulation per unit time, and *λ* represents the mean dispersal distance, conditional upon movement. For a given origin, *i*, the dispersal kernel entries, 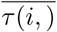, sum to 1. Computing *τ*_*ij*_ for all combinations of origins and destinations produces a transition matrix, 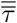. If dispersal is sex-specific, then there is a transition matrix for adult females, 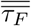, and males, 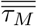. Dispersal kernels may be defined with arbitrary complexity by the user.

Movement of mosquitoes between metapopulations is computed at the end of each time step. Until this point in the document, we have been describing population dynamics at a single node; however, to accommodate all nodes on a landscape, we increase the dimensionality of our matrix/vector for the number of adult females and males having each genotype at time *t* to also include the metapopulation number. The number of adult males having each genotype in each metapopulation at time *t*, following migration, is then given by:

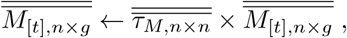

and the equivalent number for adult females, considering also the genotype of the male with whom they mated, is given by:

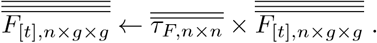

Here, the first dimension of the matrix, 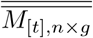, and tensor, 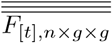, represents the metapopulation, *i ∈*{1, *…, n*}, the second dimension represents the genotype, *j ∈*{1, *…, g*}, and the third dimension (for adult females) represents the genotype of the male mate, *k ∈*{1, *…, g*}. The matrix/tensor cross product effectively sums migrating mosquitoes of a given (mated) genotype from all nodes across the network, including the origin node, based on the predictions of the dispersal kernel. The resulting numbers are used for the implementation of the inheritance and life history population dynamics in the next time step.

## 6 Stochastic simulations

Simulations in **MGDrivE** can be run either in deterministic or stochastic form, with stochasticity being able to be switched on or off in several parts of the model. In the stochastic implementation of the model:

- Daily egg production for adult females having a given genotype is Poisson-distributed, with mean equal to the number of adult females having that genotype, 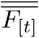, multiplied by their genotype-specific fecundity (daily egg production rate), 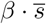.
- offspring genotypes are distributed according to a Multinomial distribution, with probabilities determined by the relevant entries of the inheritance cube, 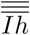, and viability mask, 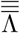, for a given mated maternal genotype.
- offspring sex is distributed according to a Binomial distribution, with probabilities determined by the offspring genotypes and their genotype-specific sex ratios, 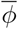.
- Female choice of male mate follows a Multinomial distribution, with probabilities of choosing each male mate determined by their mating competitiveness, 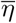, and genotype frequencies in the population, 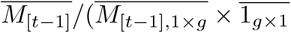.
- Daily survival at all life stages and pupatory success all follow Binomial distributions at the population level, with probabilities equal to rates provided in the population dynamic equations.
- Destination choice for migrating mosquitoes follows a Multinomial distribution, with probabilities taken from the relevant row of the metapopulation transition matrix, 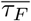 for females and 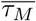 for males. An option is also available for destination choice to follow a Dirichlet distribution, the “concentration parameter” of which allows for greater variance in movement than permitted by the Multinomial distribution.

